## Supplementary Information for "Retention Time Prediction Using Neural Networks Increases Identifications in Crosslinking Mass Spectrometry"

#### Contents

**S1 Description of xiRT** **S-2**

**S2 Hyper-Parameter Optimization on Linear Data** **S-13**

**S3 xiRT Explainability Analysis** **S-14**

**S4 pLink2 Processing** **S-17**

#### List of Figures

S1 xiRT network architecture . . . S-3
S2 Cross-validation results (linear peptide data) . . . S-4
S3 Crosslink identifications over fractions / time. . . . S-4
S4 Peptide properties from entrapment and target database . . . S-5
S5 Hyper-parameter optimization for xiRT (crosslinked peptide data) . . . S-5
S6 xiRT prediction performances with single-task and multi-task set-ups . . . S-6
S7 xiRT execution times with single-task and multi-task set-ups . . . S-6
S8 Redundancy of CSMs across SCX / hSAX fractions . . . S-7
S9 Learning curves (crosslinked peptide data) . . . S-7
S10 Prediction of RP retention for Fanconi anaemia monoubiquitin ligase complex data . . . S-8
S11 SHAP explanation for a peptide observed in hSAX . . . S-9
S12 SHAP explanation for a peptide observed in SCX . . . S-9
S13 Global SHAP explanations for peptide observation in SCX / hSAX / RP . . . S-10
S14 UMAP-based embedding space visualization . . . S-11
S15 Combining pLink2 with xiRT . . . S-12

#### List of Tables

S1 First parameter grid for the optimization on linear peptide data. . . . S-14
S2 Second parameter grid for the optimization on linear peptide data. . . . S-14
S3 RT features used for prediction on *E. coli* data set. . . . S-15
S4 Unique and redundant CSMs across hSAX and SCX fractions. . . . S-16
S5 Rescoring gains with different number of chromatographic dimensions. . . . S-16

*authors contributed equally

### S1 xiRT: Multi-task Retention Time Prediction using Neural Networks

#### Overview

The schematic architecture of the xiRT was presented in Figure 1 of the manuscript, while Fig. S1 shows a more detailed view (**exemplary configuration**). Here, we want to give more details about the individual layers. The input layer dimension is dynamically defined by the longest peptide that was identified in the set of PSMs/CSMs. In the example in Fig. S1 this was set to *59*. Subsequently, the input is passed to a predefined embedding layer in TensorFlow. The embedding layer finds a continuous vector representation from a list of positive integers. A hyper-parameter for the network is the length of the embedding vector, here set to *50*.

#### Siamese Architecture

The heart of the xiRT network is a recurrent layer where we either used a Gated Recurrent Unit (GRU-) [2] or a Long short-term memory (LSTM)-layer [3]. These layers are especially suited for the handling sequential data, e.g. language data or peptide sequences. They are available as GPU and CPU implementations in TensorFlow and can thus be used interchangeably within xiRT. The central assumption for recurrent layers is that the order of the input (here amino acids) plays a pivotal role in the prediction process [4]. By optionally applying a bidirectional GRU/LSTM layer, the input sequence is handled forward and backward. To speed up the training process, the activations are further batch-normalized to $\mu = 0, \sigma = 1$. The above-described parts of the network are designed in a Siamese fashion. That means that two input sequences (i.e. the individual peptides in a crosslink) are passed to their custom inputs. However, these layers process the input in the same manner since they share the same weights. The combination of the outputs from the Siamese network can be handled in multiple ways. In Fig. S1 an *Add-layer* was used, which simply adds the two inputs element-wise. For the retention time prediction of linear peptides there is only a single input and thus no Siamese or additive layers are necessary.

#### Task Specific Layers

The architecture described above is also shared between the different prediction tasks. In this manuscript, we developed a multi-task network that predicts peptide retention behaviour during SCX, hSAX and RP chromatography. For this, the individual task-networks were designed in a symmetric fashion. They are defined by a sequence of layers with: $layer_i$ = *Dropout(BatchNormalization(Dense(x)))*. Per default we used $i = 3$ and a pyramid-like structure for the dense layers with $n_{neurons} = [300, 150, 75]$. The default dropout-rate was set to 0.1 for all dense layers. Moderate kernel regularization (l2, $\lambda = 0.001$) was also used.

For each task, a custom prediction layer and a loss function are defined. The two employed fractionation techniques SCX and hSAX are handled as ordinal regression problems in which sigmoid activations were used and binary cross-entropy as loss. For the RP, we used a linear activation function and the mean squared error as loss function. Note that the handling of data from fractionation also allows to treat the problem as classification or as regression task and thus the use of softmax or linear activation functions are possible (also configurable in xiRT). The total loss is computed as weighted sum of the three individual losses, e.g. $loss_{total} = w_{fractionation} * (loss_{SCX} + loss_{hSAX}) + loss_{RP}$. Using *Adam* (Adaptive Moment Estimation) as optimizer, the learning rate was fixed to 0.001 during development and optimization on linear data. After optimization for crosslink data a higher learning rate (0.01) achieved faster convergence with similar accuracy together with a batch-size of 256 and was therefore chosen as default value in xiRT.

#### Implementation Details

xiRT is implemented in the popular deep learning framework TensorFlow 2.0 [5]. All training and prediction scripts were run on a TITAN X (Pascal) with 12.8 GB of memory. The usage of a dedicated GPU allows to use optimized recurrent layers in TensorFlow. These layers have a "CuDNN"-prefix, e.g. CuDNNGRU. CuDNNLSTM. Our implementation can also be used on systems without GPUs, at the cost of higher run time. TensorFlow also allows the usage of so-called callbacks. The most important callbacks in our implementation are 1) *ReduceLRonPlateau*, 2) *EarlyStopping* and 3) *ModelCheckpoint*. The 1) callback is used to reduce the learning rate by a factor of 0.5 when the performance has not improved in 15 epochs by a minimum delta of 1e-4. Similarly, 2) is used to speed up the training by stopping the process if no improvements were achieved in a configurable number of epochs. In addition, for the final model the best weights over all epochs are chosen based on the performance on the validation split. Finally, 3) is used to store the weights and model architecture on disk. This allows applying the best model for the respective cross-validation folds and the candidate rescoring. For transfer learning applications these trained models can also be used for new data sets. Most parameters of the network such as learning rate, optimizer, batch size, epochs, callback settings, number of layers / neurons can be adapted through a dedicated YAML file. The online documentation for xiRT on GitHub contains examples for various training and RT dimension scenarios.

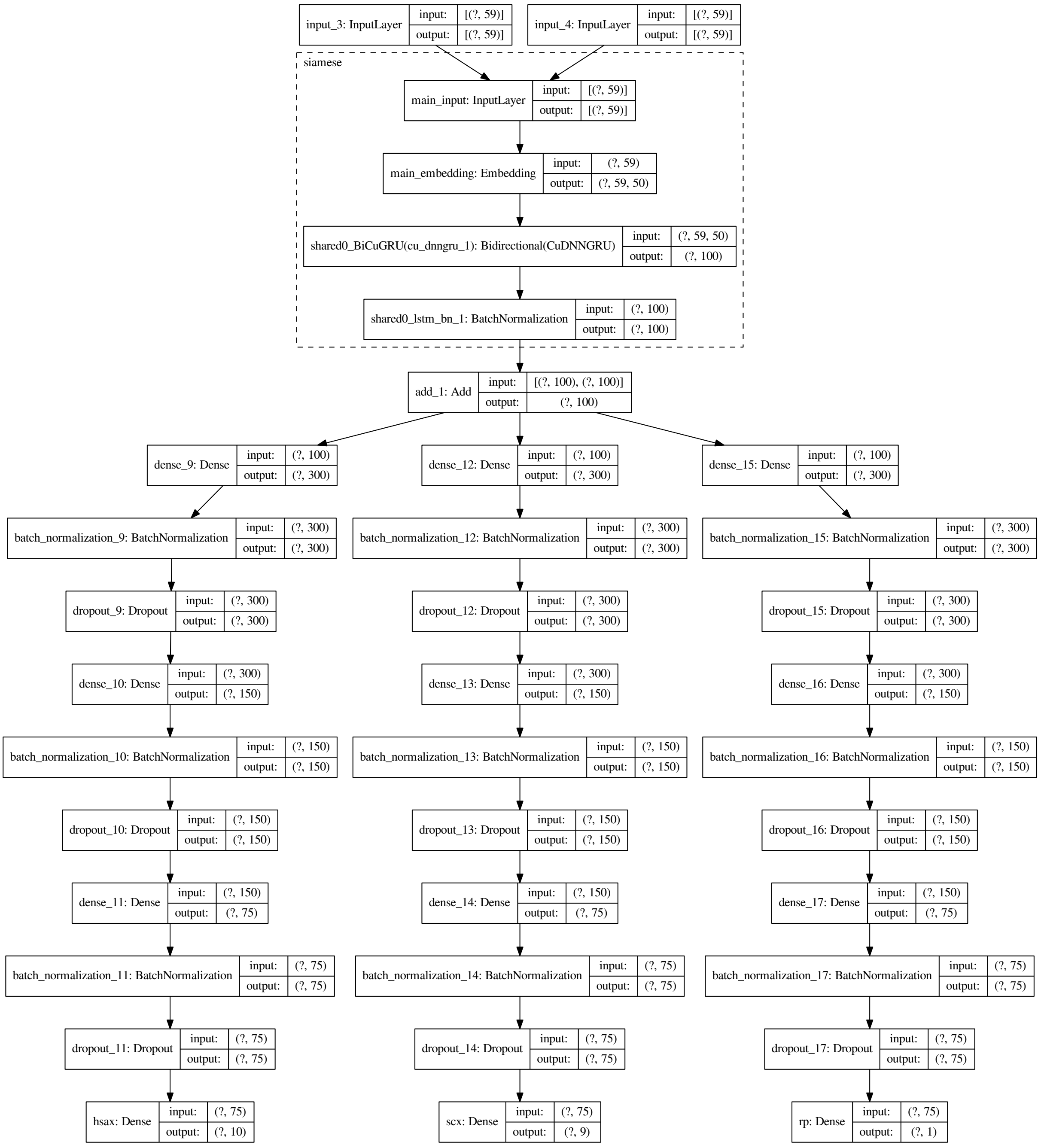


Figure S1: Example parameterization of xiRT. Dashed box represents the Siamese network part. Boxes represent individual layers with their names, input and output dimensions. Question marks represent the unknown batch-size at compilation time of the network.

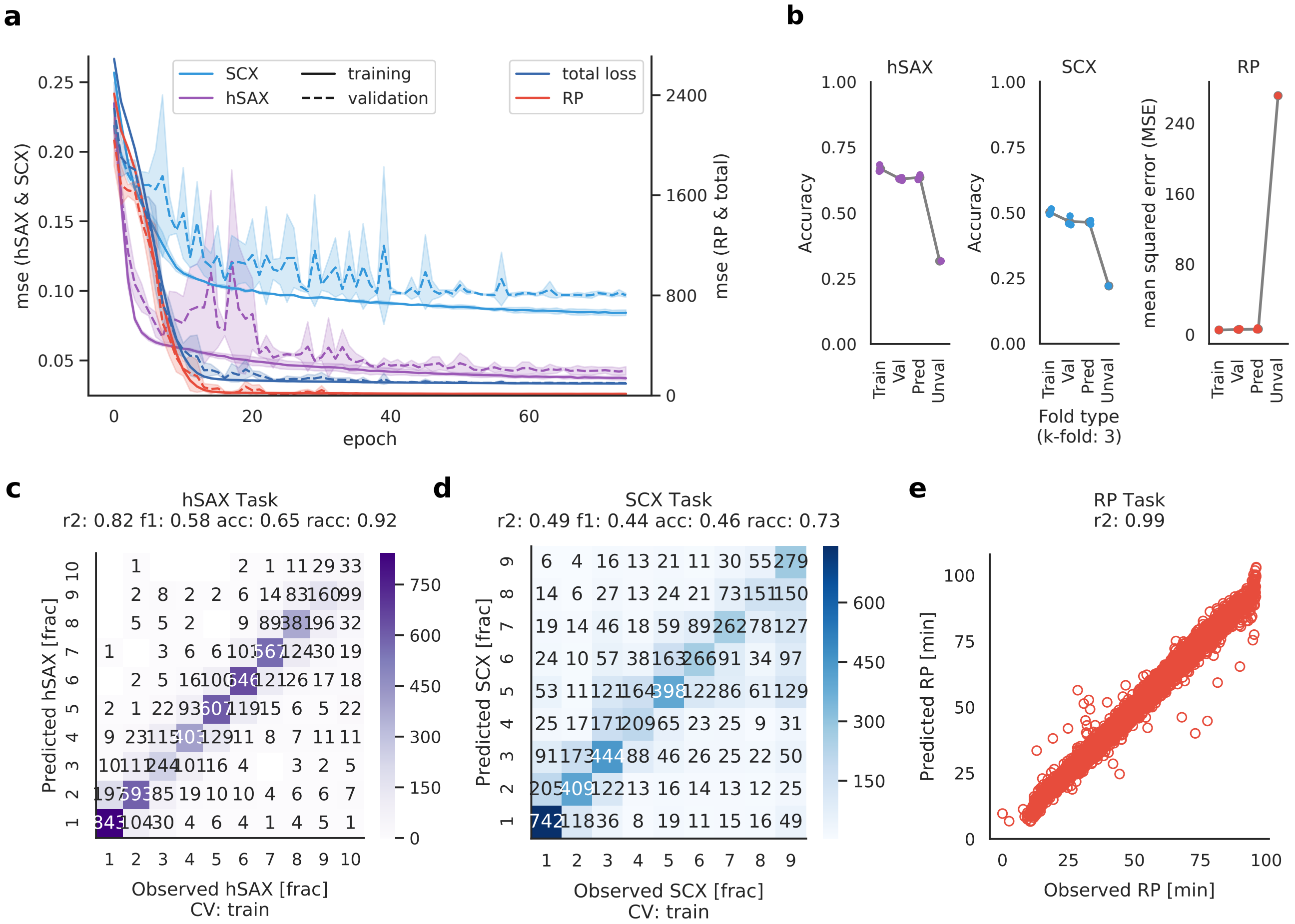


Figure S2: Cross-validation results from applying xiRT on linear peptide input. a) Training performance on all tasks over 75 epochs. Confidence intervals show standard deviation from a 3-fold CV. b) Evaluation metrics for all tasks on the different CV folds. c-e) Representative results for a random prediction fold. Abbreviations: val, validation; pred, prediction, unval, unvalidated; mse, mean squared error; acc, accuracy.; racc, relaxed accuracy ($|error| \leq 1$).

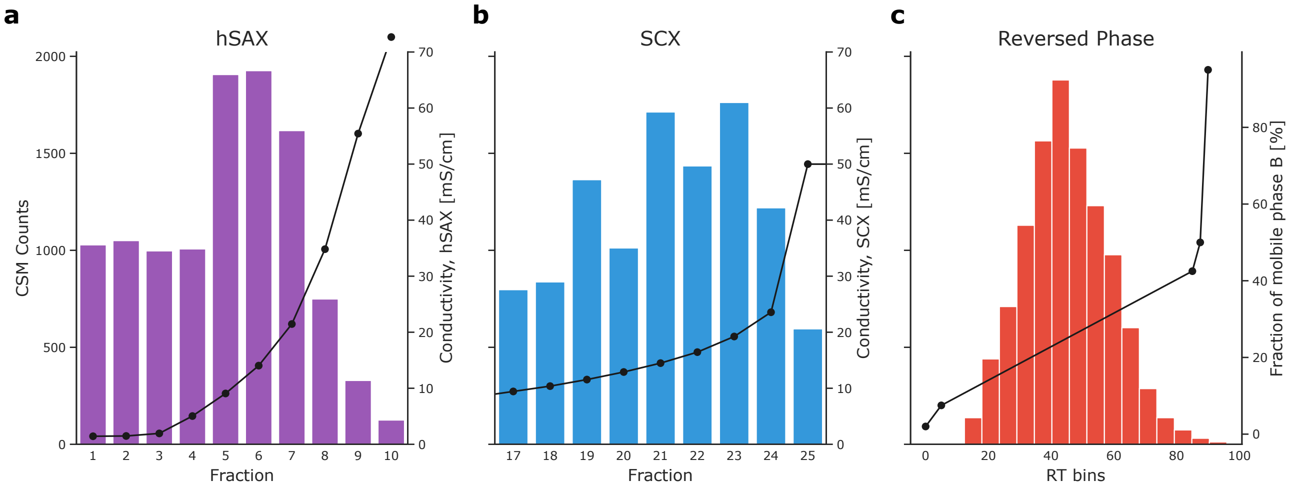


Figure S3: Crosslink identifications over fractions / time. (a-b) Distribution of CSMs across native fraction numbers from the off-line fractionation based on strong cation exchange (SCX) and hydrophilic strong anion exchange (hSAX) chromatography. Black lines indicate the eluent concentration (represented by conductivity) at the beginning of the fraction. (c) Distribution of CSMs across reversed-phase retention time bins. Black lines indicate the eluent concentration (fraction of eluting mobile phase B) at the beginning of the fraction. Data corresponds to 11055 CSMs at 1% FDR, all target-target hits excluding matches involving human proteins.

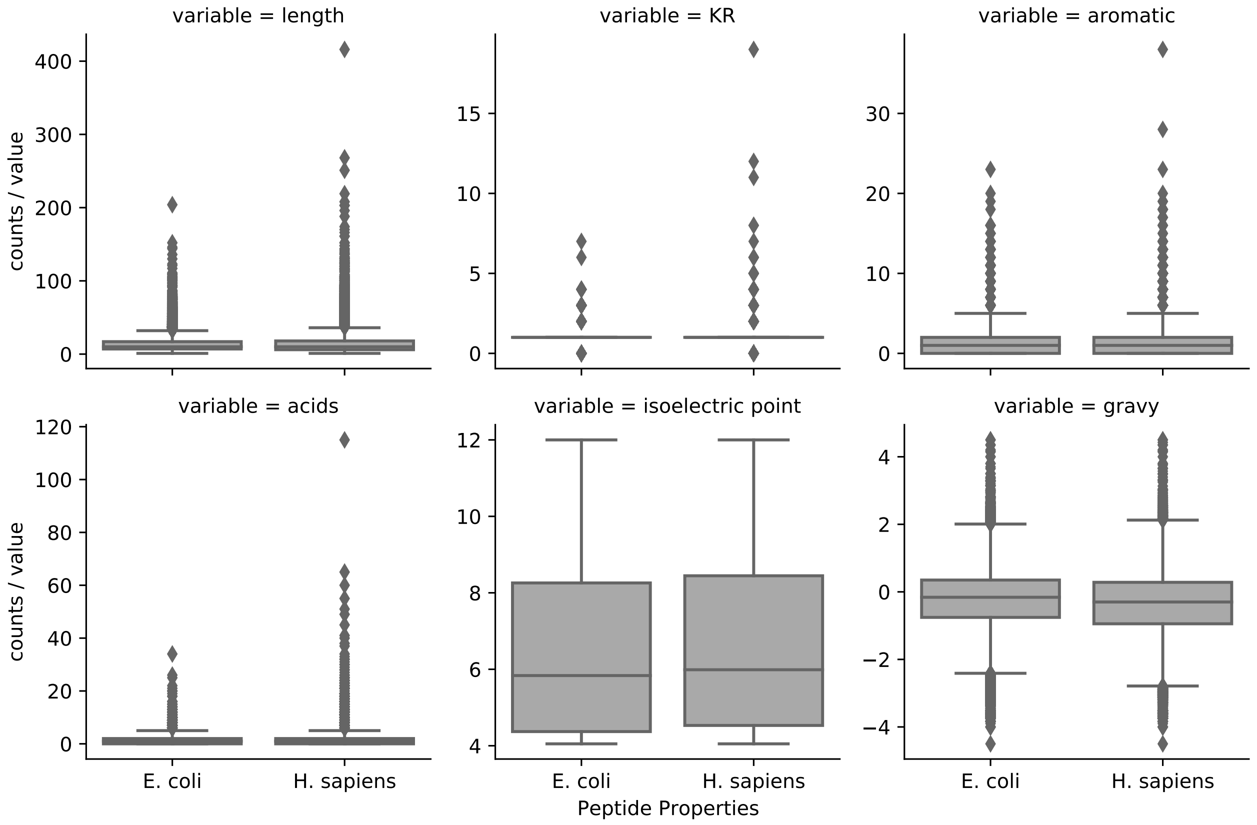


Figure S4: Comparison of peptide properties from the E. coli target database and the H. sapiens entrapment database. Variables: KR, K/R count in peptide; aromatic, F/Y/W counts; acids, D/E counts; isoelectric point and GRAVY were computed using Biopython [1]. Boxplots show the median as line and the whiskers show the 1.5x interquartile range, points represent the outliers.

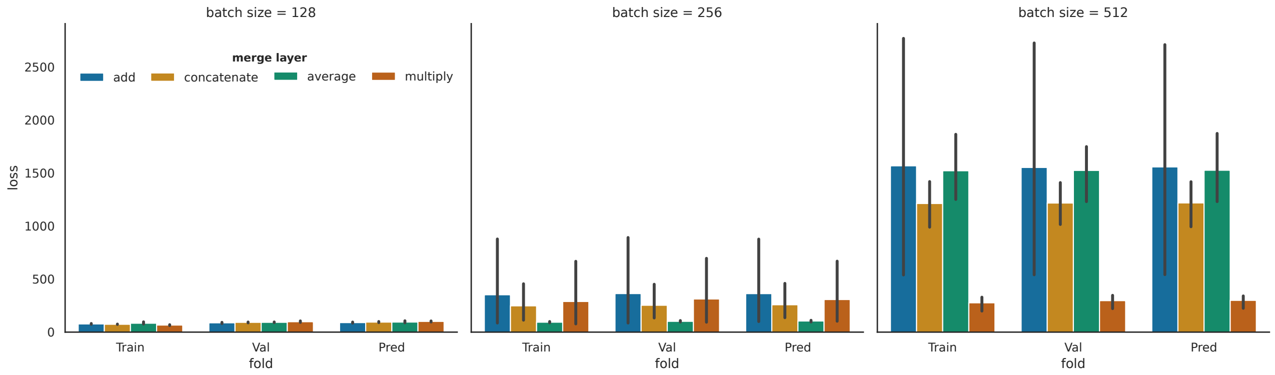


Figure S5: Hyper-parameter optimization for xiRT. Appropriate hyper-parameters were assessed following cross-validation on crosslinked peptides. Bars are sorted by the lowest loss value in the prediction folds. Error bars represent the standard deviation.

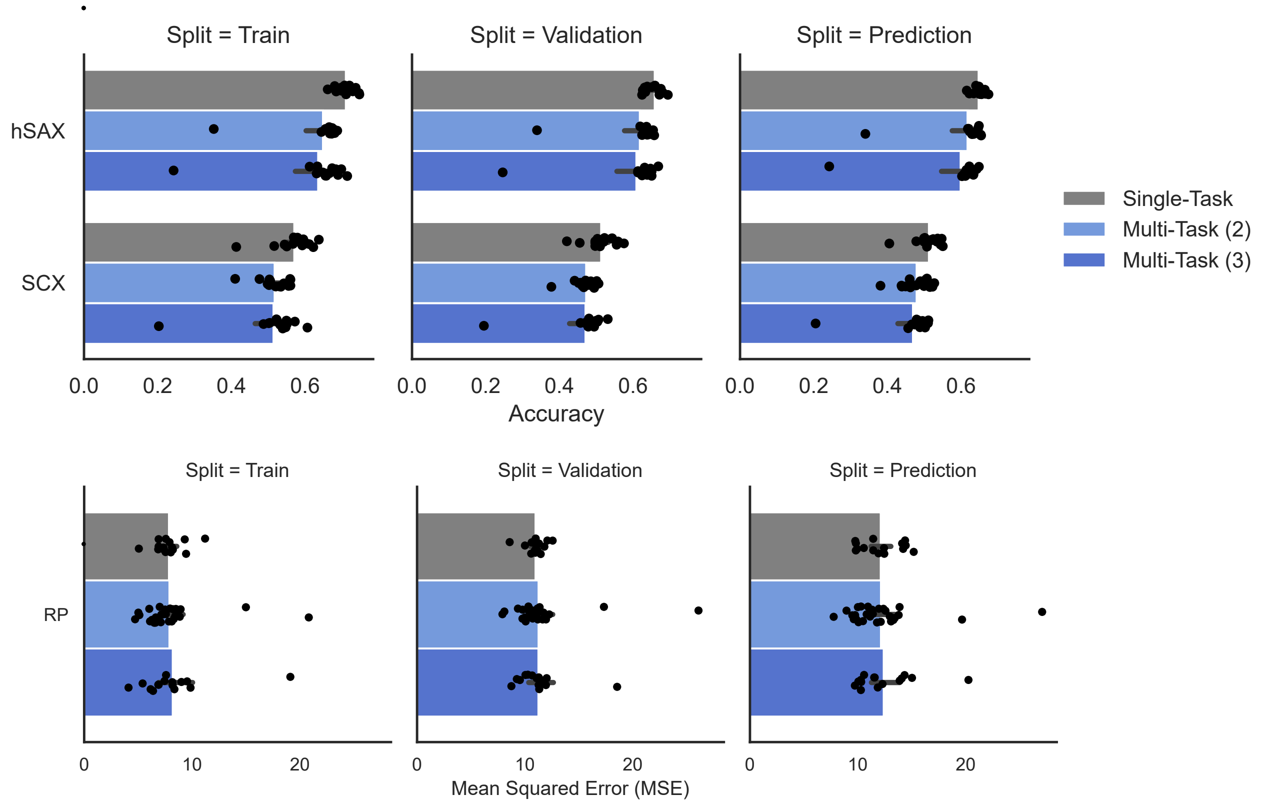


Figure S6: xiRT performance for single- and multi-task parameterizations (with either (2) or (3) tasks). Bars show the mean value from five replica of 3-fold CV (every dot represents a single CV result). ANOVA (type 2) results fail to reject the null hypothesis of an equal mean in all groups at $\alpha = 0.05$ (prediction split only). Results were: SCX ($F = 2.7$, $p - value = 0.08$), hSAX ($F = 1.69$, $p - value = 0.20$), RP ($F = 0.04$, $p - value = 0.96$), with 2 degrees of freedom and 42 total df for the SCX/hSAX test and 57 total df for the RP test.

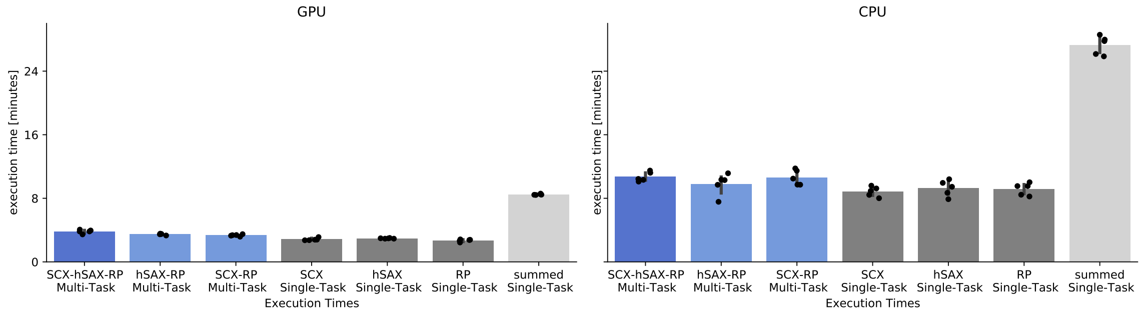


Figure S7: xiRT benchmark for single- and multi-task parameterizations on different hardware. Every parameter was tested in five replicates. For the 'summed' single-task estimate random replicates were paired. GPU analysis was performed on Intel(R) Xeon(R) CPU E5-1620 v4 @ 3.50 GHz equipped with an TITAN X (Pascal) with 12GB memory. CPU analysis was performed on Intel(R) Core i7 6700K CPU @ 4.00 GHz, with 32GB DDR4 memory.

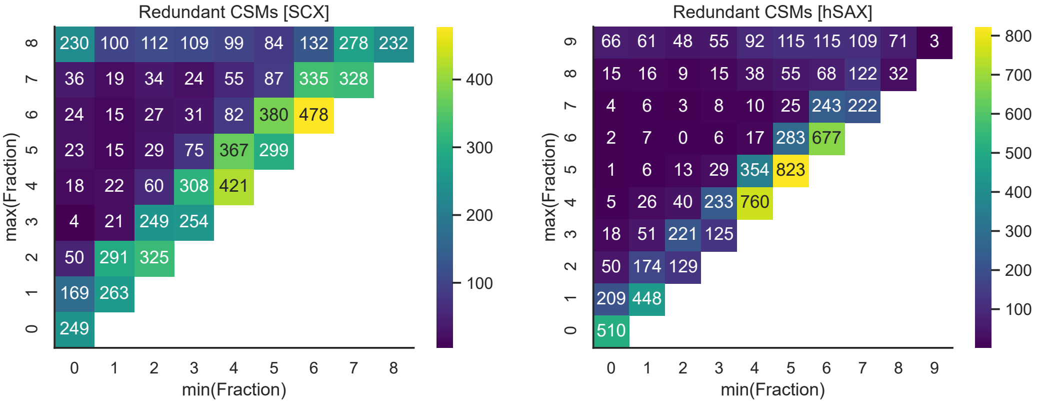


Figure S8: Redundancy of CSMs across SCX / hSAX fractions (6843 non-unique CSMs at at 1% FDR). If a CSM was identified in multiple fractions, the span (min and max of the fraction) was calculated and visualized. For example, if a CSM was identified in the fractions 2,3,4, the span would be (2, 4), leading to an increase in the plot at x=2 and y=4. In other words, 60 CSMs were indeed observed with a span of (2, 4). CSM redundancy was high in the last fractions (8 for SCX, 9 for hSAX) emphasizing ambiguous retention behavior. Fraction numbers were transformed to start at zero.

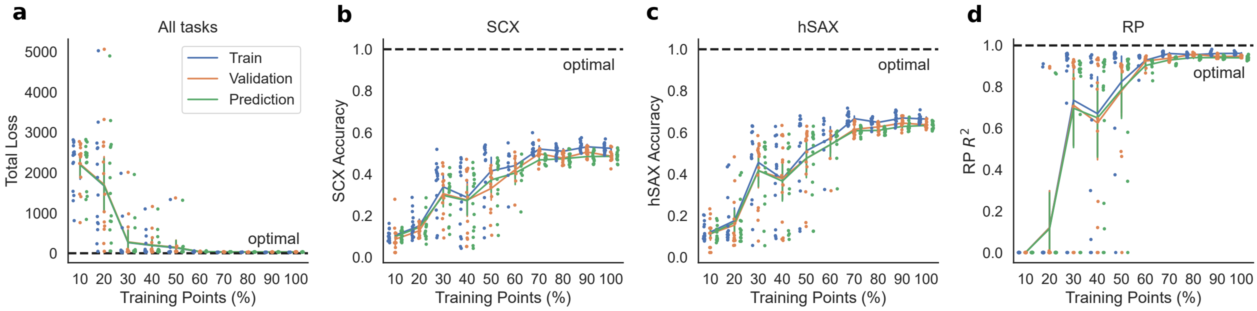


Figure S9: Learning results on crosslink data from 3-fold cross-validation (five replicates). a) total unweighted loss is shown, b and c) classification accuracy and d) $R^2$ for the RP prediction is shown. Data used (train, validation, prediction): 10% (387, 43, 215); 20% (774, 87, 430); 30% (1161, 130, 645); 40% (1548, 173, 860); 50% (1935, 216, 1075); 60% (2323, 259, 1290); 70% (2710, 302, 1505); 80% (3097, 345, 1720); 90% (3484, 388, 1936); 100% (3871, 431, 2151). Vertical bars show the standard deviation.

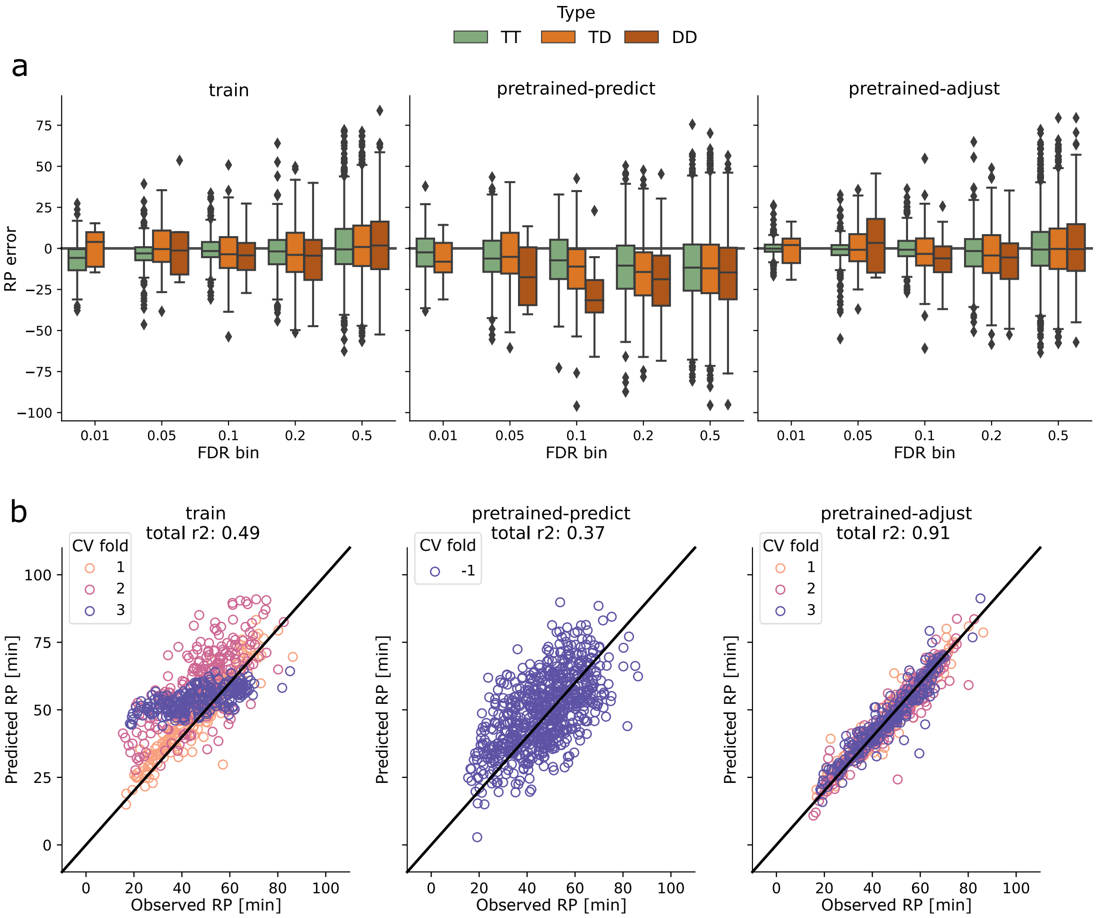


Figure S10: Reversed-phase RT prediction for the Fanconi anaemia monoubiquitin ligase complex data set[9] (FA-complex). Upper panel (a) shows the RP error for TT, TD, DDs in different FDR bins, e.g. 0.05 contains all CSMs with an $FDR \geq 0.01$ but $FDR < 0.05$. The modes *train* (solely train on FA-complex data), *pretrained-predict* (apply pretrained model, using the subset of E. coli DSS-crosslink data at 1% CSM-FDR without fine-tuning), *pretrained-adjust* (load the previously described model, include fine-tuning the network during cross-validation). Boxplots follow the description from Fig. S4. Lower panel (b) shows the individual cross-validation predictions for the three training set-ups. For the pretrained-predict model, no cross-validation was performed. While the data contained  1400 CSMs at 1% CSM-FDR, only about 700 CSMs were used by xiRT due to high peptide sequence redundancy.

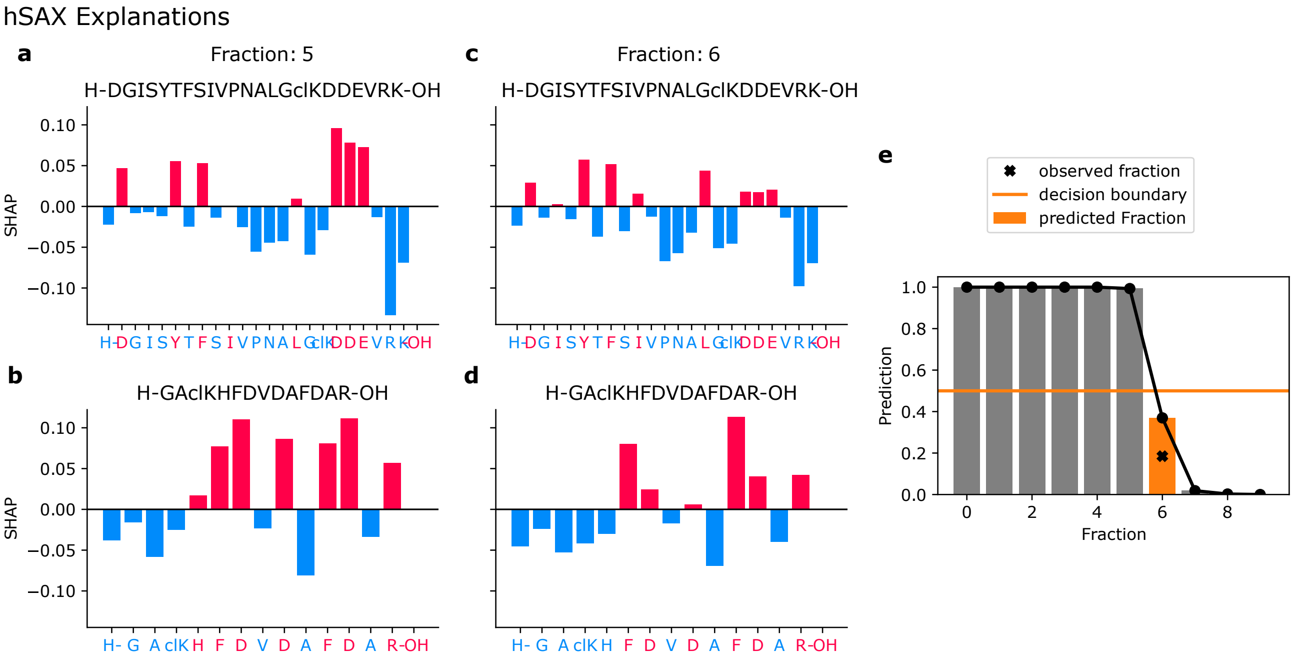


Figure S11: SHAP explanations for the hSAX elution behavior of a crosslinked peptide that eluted in hSAX fraction 6 (0-based). a-b) SHAP values for the crosslinked peptide DGISYTFSIVPNALGkDDEVRK-GAkHFDVDAFDAR (H- = N-terminus, clK = crosslinked lysine residue, -OH = C-terminus) for the prediction to elute in fraction 5. c-d) SHAP values for the peptide to elute in fraction 6. e) Predicted output of the network (ordinal fraction prediction) that is translated into the predicted fractions. The fraction is determined by the first prediction that yields a probability lower than 0.5 (orange line). Negative (blue) SHAP-values contribute towards an earlier elution, while positive (red) SHAP-values contribute towards later elution.

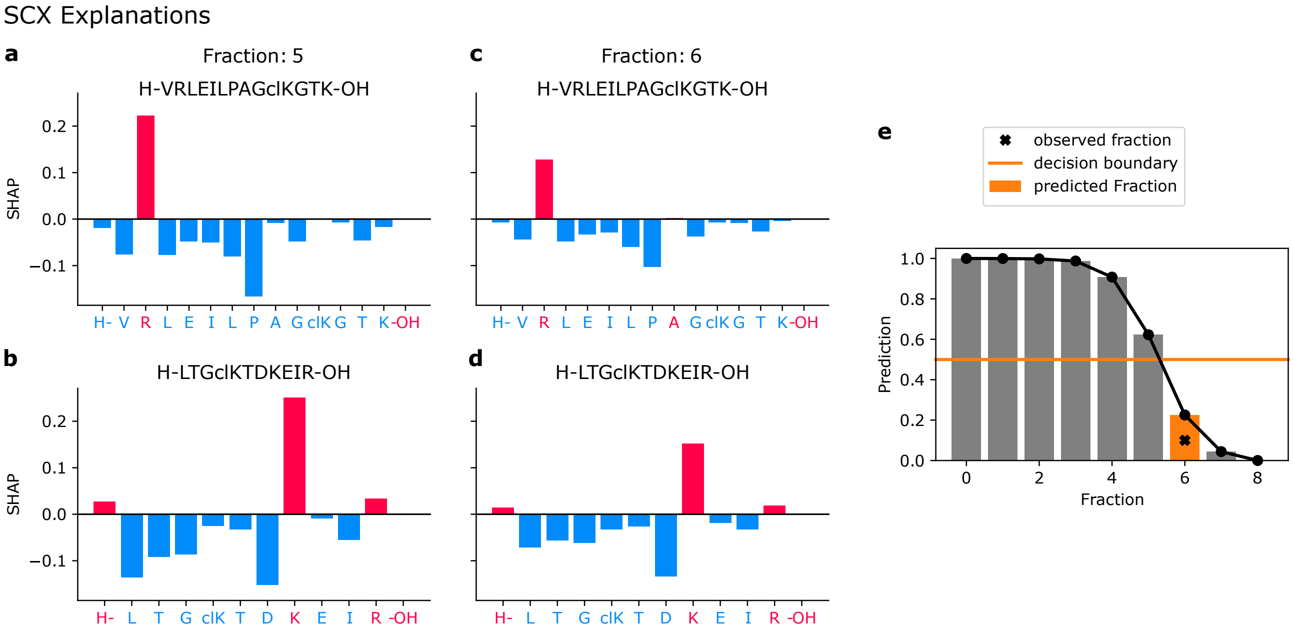


Figure S12: SHAP explanations for the SCX elution time of a crosslinked peptide that eluted in SCX fraction 6 (0-based). See Fig. S11 for a detailed description.

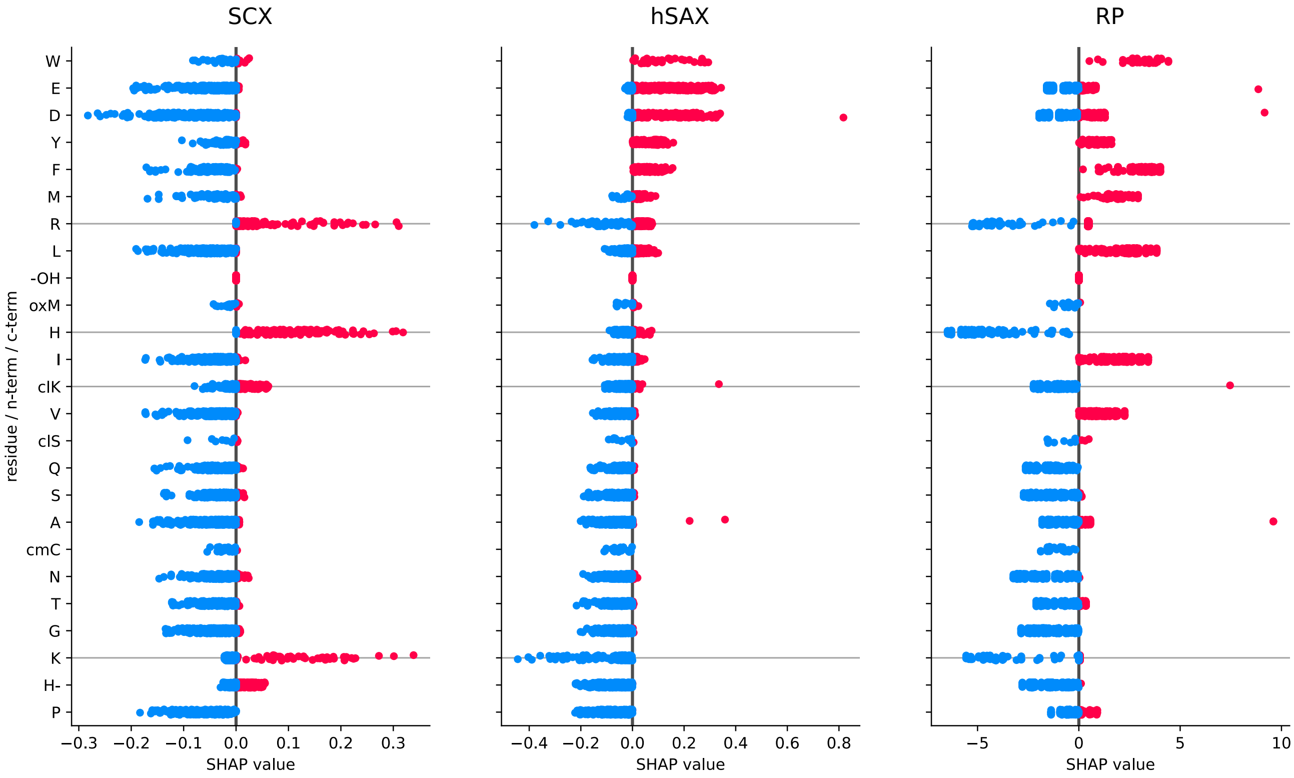


Figure S13: SHAP values for individual amino acids of crosslinked peptides contributing towards early or late elution in SCX/hSAX/RP separations. Positive (red) SHAP values contribute towards later elution, negative (blue) SHAP values contribute towards earlier elution. Horizontal grey lines highlight R, K, clK, H residues. SHAP values were computed for 200 randomly drawn CSMs passing 1% CSM-FDR.

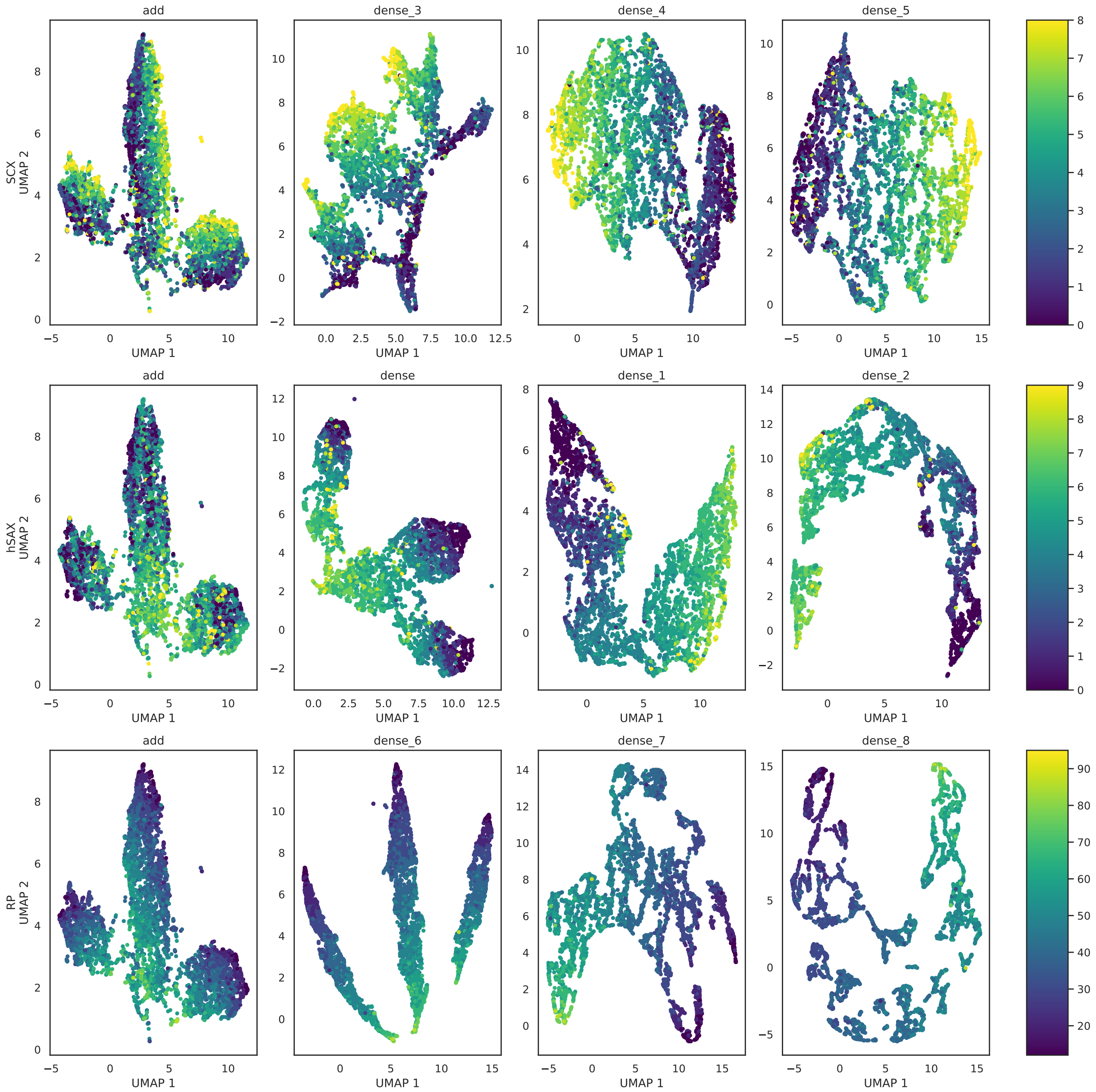


Figure S14: Visualization of the embedding space throughout the network for each task. UMAP parameters: metric, Euclidean; min_dist, 0.0; n_neighbors, 15. Individual plots correspond to the add (shared) and dense (task-specific) layers in Fig. S1. UMAP was applied to a cross-validation model from the E. coli DSS data set. Note that the given parameterization of UMAP might be suboptimal for all the selected embedding spaces.

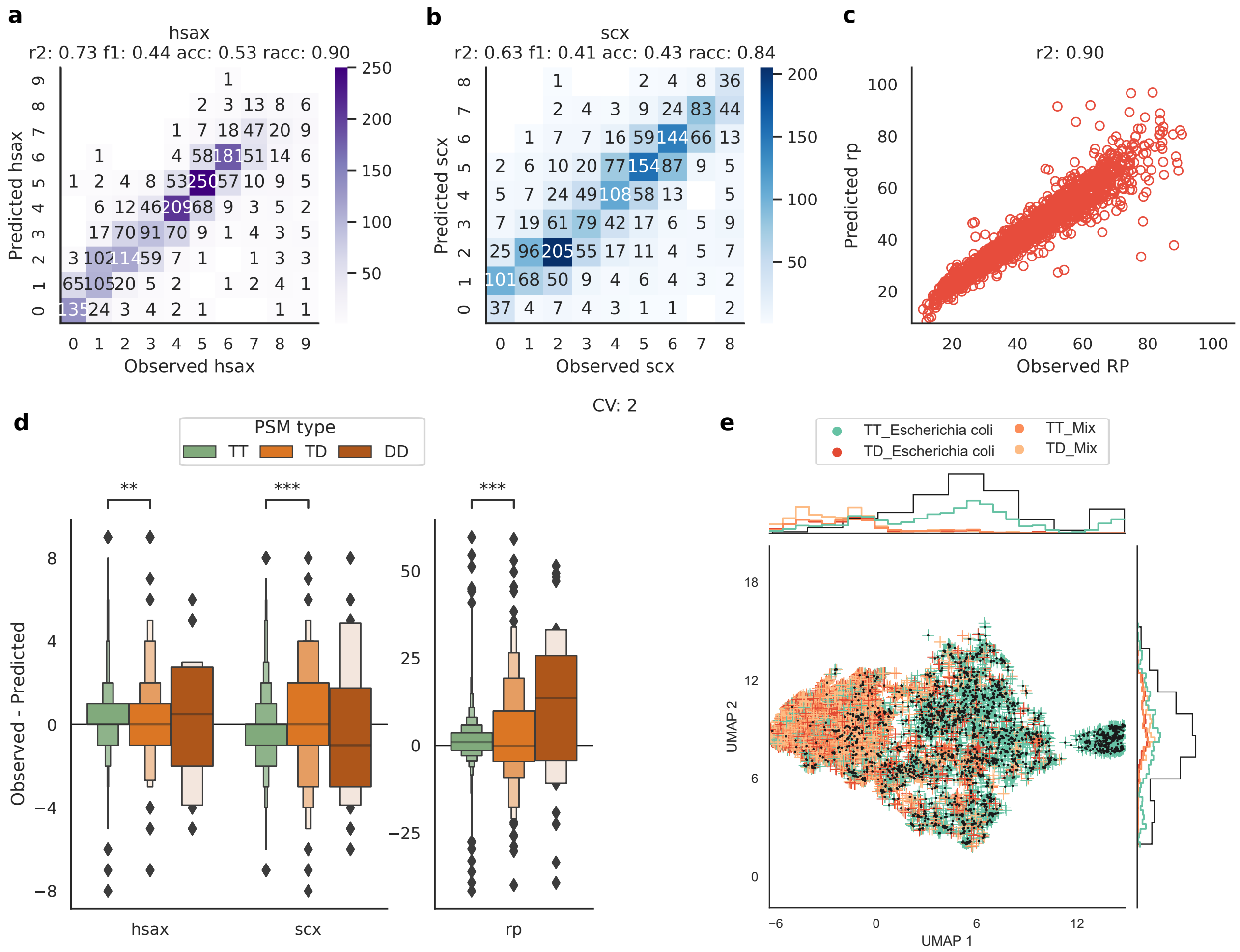


Figure S15: Evaluation of combining xiRT with pLink2 search results. a-c) Cross-validation results on the second prediction fold using all three RT dimensions (hSAX, SCX, RP). pLink2 data was filtered to a Q-value of 0.01 for the training process in xiRT. d) Error characteristics for TT (n=6436), TD (n=214) and DD (n=34). P-values are derived from a two-sided, independent t-test with Bonferroni correction between TT and TD observations. P-values: 1.545e-03 (hsax), 5.667e-04 (scx), 3.381e-04 (rp), test-statistics: -3.364e+00 (hsax), -3.632e+00 (scx), -3.586e+00 (rp). e) Dimensionality reduced feature space using UMAP with default parameters. Black dots represent CSMs that passed the 0.01 Q-value cutoff. Only heteromeric crosslink spectrum matches are shown.

### S2 Hyper-Parameter Optimization on Linear Data

Neural networks are subject to many parameters that need tuning to achieve the best possible performance. Based on our initial work on the prediction of hSAX RTs for linear peptides [6], we came up with an initial architecture and then optimized it manually. Further, we choose a step-wise approach to find suitable hyper-parameters during a 3-fold cross-validation search. The first grid of hyper-parameters is shown in Table S1. For the CV, we split the data into a *training* fold, a *validation* fold (10% of the training fold data) and a testing fold per CV iteration. We also use the term *prediction fold* synonymous to the *testing fold* since we only use the testing fold predictions for CSM rescoring later on. All decoy-PSMs and identifications with a FDR higher than the selected training FDR are assigned to an *unvalidation* fold. After the first round of CV on a set of 576 parameters, another grid-search (320 parameters) was performed with adjusted hyper-parameters (Table S2). This second grid was based on the best performing parameters in the first iteration with slight variations. Note, the linear peptide identifications at 1% PSM-FDR were used for this procedure (n=20802 unique sequences, ignoring identifications to the entrapment database). For the execution of the hyper-parameter search, we again designed a snakemake [7] workflow that can run an arbitrary number of configuration files. The best final parameters were then chosen based on the means of the loss, $r^2_{rp}$, $accuracy_{hSAX}$ and $accuracy_{SCX}$ in the testing sets during CV. Note that the 2-step optimization offers a reasonable trade-off between finding optimal parameters and decreasing the necessary run time.

The best parameters from optimization (Tab. S1, Tab. S2), showed an average ($\pm$ standard deviation) R2 of 0.99 $\pm$ 0.003 for the RP task, average accuracy 64% $\pm$ 0.9 for hSAX task and 46% $\pm$ 0.7 for the SCX task (Fig. S2). By using a relaxed accuracy metric (absolute prediction error $\leq$ 1 fraction), hSAX RT prediction achieved 92% $\pm$ 0.3 and SCX RT prediction 74% $\pm$ 0.7.

The network performance across the individual CV-folds of the best parameter was very comparable in terms of training time and performance (Fig. S2a). The CV was performed on 20802 unique CSMs (train: 12481, validation: 1387, prediction: 6934 observations). The learning trajectory of the number of epochs follows a very smooth learning curve and shows a constant improvement in the training and validation fold with a small gap between the training and validation performance. We also observed that the prediction accuracy for hSAX is better than for SCX in both, training and validation data. This trend is also observable in the prediction folds (Fig. S2b). In addition, the performance drop from the validation fold to the prediction fold is rather small which is desirable and shows good generalization ability of the network. A lower prediction performance on the unvalidation split can be expected and hence serves as another quality check. The predictions were made with the best classifier from the CV split. The individual predictions for a single CV-fold are more accurate for the RP than for SCX or hSAX (Fig. S2c-e). While the RP predictions achieve an $r^2$ of 0.99, the accuracy in SCX and hSAX is limited to 0.64 and 0.45, respectively. The different behaviour of hSAX and SCX might be explained through deviating peptide separation behaviour with the applied gradients (Fig. S3). While the shape of the gradients is similar, the hSAX gradient led to a more uniform distribution of crosslinked peptides across the elution window in contrast to the more confined elution of crosslinked peptides in later SCX fractions. Therefore, adjacent SCX fractions are expected to show a higher overlap in their identifications than fractions from hSAX.

Table S1: First parameter grid for the optimization on linear peptide data.

| Parameter | Parameter Grid | Selected Parameter |
|---|---|---|
| recurrent type | CuDNNGRU | CuDNNGRU |
| recurrent units | 25, 75, 125 | 75 |
| recurrent activity l2 lambda | 0, 0.001 | 0.001 |
| recurrent kernel l2 lambda | 0 | 0 |
| dense layers | 3 | 3 |
| dense neurons | (300, 150, 75), (150, 100, 50) | (150, 100, 50) |
| dense kernel l2 lambda | (0.001, 0.001, 0.001) | (0.001, 0.001, 0.001) |
| dense dropout | (0.3, 0.3, 0.3), (0.1, 0.1, 0.1) | (0.1, 0.1, 0.1) |
| dense activation | (relu, relu, relu), (swish, swish, swish) | (swish, swish, swish) |
| embedding length | 50, 100 | 50 |
| batch size | 256, 512 | 256 |
| class weight | 1, 250, 500 | 250 |

*Note*: Parameters with a prefix "recurrent" were used for a single recurrent layer. Parameters with the "dense" prefix were used for the task specific layers. A total set of 576 parameter combinations were used during the grid search. Remaining settings were left at defaults. The best parameter was determined based on the testing folds in a 3-fold CV experiment. Training time was limited to 75 epochs and early stopping patience was set to 15.

Table S2: Second parameter grid for the optimization on linear peptide data.

| Parameter | Parameter Grid | Selected Parameter |
|---|---|---|
| recurrent type | CuDNNGRU, CuDNNLSTM | CuDNNGRU |
| recurrent units | 75 | 75 |
| recurrent activity l2 lambda | 0, 0.001 | 0.001 |
| recurrent kernel l2 lambda | 0, 0.001 | 0.001 |
| dense layers | 3 | 3 |
| dense neurons | (300, 150, 75) | (300, 150, 75) |
| dense kernel l2 lambda | (0.001, 0.001, 0.001) | (0.001, 0.001, 0.001) |
| dense dropout | (0.2, 0.2, 0.2), (0.1, 0.1, 0.1 ) | (0.1, 0.1, 0.1) |
| dense activation | (relu, relu, relu), (swish, swish, swish) | (relu, relu, relu) |
| embedding length | 50, 75 | 50 |
| batch size | 256 | 256 |
| class weight | 1, 50, 100, 200, 250 | 200 |

### S3 xiRT Explainability Analysis

In this section we describe the analysis of the SHAP values from the learned multi-task model. For this, the used tensorflow-version needed to be downgraded to 1.15 together with SHAP (v. 0.36.0). As background data 100 randomly chosen CSMs were provided. To use the DeepExplainer, the trained network had to be dissected into the single tasks. Furthermore, the ordinal regressions setup for hSAX and SCX complicates the analysis since each sigmoid activation of the output vector can be explained via SHAP (padded positions were ignored). Therefore, we only focused on the SHAP values for the relevant prediction decision, i.e. the sigmoid activiation that was <= 0.5. With this special model architecture, the returned SHAP-values failed the 'check_additivity' flag in the SHAP package and the check was thus disabled. However, the magnitude and overall explanations from the DeepExplainer show realistic feature importance values for the RT contributions on residue level. Since the SHAP values only represent an approximation of the contributions we further explored their magnitude. In Fig. S11 we demonstrate the explainability via SHAP of a crosslinked peptide's predicted retention time (hSAX fraction). The residues D, E, R and K behave mostly as expected. In addition, aromatics (Y, F) also contribute to stronger retention and hence later elution times, while A contributes towards earlier elution times. These observations are in line with an earlier study on the hSAX RT behavior[6]. Similarly, an explanation for a SCX prediction is shown in Fig. S12. The global SHAP values based on the *raw sequence* inputs to xiRT are shown in Fig. S13. For hSAX again, D, E, F, Y, W belong to the major contributors towards extended retention times. For SCX, the positive contribution is mainly attributed towards R, K and H. Note that crosslinked K residues, contribute much less towards later elution times than non-crosslinked K residues.

Table S3: RT features used for prediction on *E. coli* data set.

| # | Feature Name | Description |
|---|---|---|
| 1 | hsax-error | crosslinked - raw error between observed and predicted (hSAX) |
| 2 | scx-error | crosslinked - raw error between observed and predicted (SCX) |
| 3 | rp-error | crosslinked - raw error between observed and predicted (RP) |
| 4 | hsax-error-peptide1 | raw peptide 1 error (hSAX) |
| 5 | scx-error-peptide1 | raw peptide 1 error (SCX) |
| 6 | rp-error-peptide1 | raw peptide 1 error (RP) |
| 7 | hsax-error-peptide2 | raw peptide 2 error (hSAX) |
| 8 | scx-error-peptide2 | raw peptide 2 error (SCX) |
| 9 | rp-error-peptide2 | raw peptide 2 error (RP) |
| 10 | peptide1_mean | median of all peptide1 error (absolute values) |
| 11 | peptide1_sum | sum of all peptide1 error (absolute values) |
| 12 | peptide1_max | maximum of all crosslinked errors |
| 13 | peptide1_min | minimum of all peptide1 errors |
| 14 | peptide2_mean | median of all peptide2 errors (absolute values) |
| 15 | peptide2_sum | sum of all peptide2 errors (absolute values) |
| 16 | peptide2_max | maximum of all peptide2 errors |
| 17 | peptide2_min | minimum of all peptide2 errors |
| 18 | cl_mean | median of all crosslinked errors (absolute values) |
| 19 | cl_sum | sum of all crosslinked errors (absolute values) |
| 20 | cl_max | maximum of all crosslinked errors |
| 21 | cl_min | minimum of all crosslinked errors |
| 22 | initial_prod | log2 product (absolute values + 0.1) of all initial errors (#1-9) |
| 23 | initial_sum | sum (absolute values) of all initial errors (#1-9) |
| 24 | initial_min | minimum (absolute values) of all initial errors (#1-9) |
| 25 | initial_max | maximum (absolute values) of all initial errors (#1-9) |
| 26 | hsax-error_square | squared hsax-error for crosslinked errors |
| 27 | hsax-error_abs | absolute hsax-error for crosslinked errors |
| 28 | scx-error_square | squared scx-error for crosslinked errors |
| 29 | scx-error_abs | absolute scx-error for crosslinked errors |
| 30 | rp-error_square | squared rp-error for crosslinked errors |
| 31 | rp-error_abs | absolute rp-error for crosslinked errors |
| 32 | hsax-error-peptide1_square | squared hsax-error for peptide1 errors |
| 33 | hsax-error-peptide1_abs | absolute hsax-error for peptide1 errors |
| 34 | scx-error-peptide1_square | squared scx-error for peptide1 errors |
| 35 | scx-error-peptide1_abs | absolute scx-error for peptide1 errors |
| 36 | rp-error-peptide1_square | squared rp-error for peptide1 errors |
| 37 | rp-error-peptide1_abs | absolute rp-error for peptide1 errors |
| 38 | hsax-error-peptide2_square | squared hsax-error for peptide2 errors |
| 39 | hsax-error-peptide2_abs | absolute hsax-error for peptide2 errors |
| 40 | scx-error-peptide2_square | squared scx-error for peptide2 errors |
| 41 | scx-error-peptide2_abs | absolute scx-error for peptide2 errors |
| 42 | rp-error-peptide2_square | squared rp-error for peptide2 errors |
| 43 | rp-error-peptide2_abs | absolute rp-error for peptide1 errors |

*Note*: Features computed from xiRT predictions. All errors or predictions are derived from the same xiRT model for crosslinked peptides. In the case of individual peptide predictions (peptide1/peptide2), the second sequence in the input is set to all-zeroes. This feature set was used in the *E. Coli* analysis with all three RT dimensions.

Table S4: Unique and redundant CSMs across hSAX and SCX fractions.

| | Total | Unique | Redundant | hSAX (red/same) | hSAX (red/diff) | SCX (red/same) | SCX (red/diff) |
|---|---|---|---|---|---|---|---|
| Counts | 39226 | 4500 | 6843 | 3729 | 3114 | 2849 | 3994 |
| % | 100 | 40 | 60 | 33 | 27 | 25 | 35 |

*Note*: Unique and redundant CSM identifications at 1% CSM-FDR (separate for heteromeric and self-links). Unique CSMs are combinations of peptide 1, peptide 2, link site and charge state that were only identified once. Redundant (red) CSMs were identified more than once and thus can either have different RT times ("diff") or the same RT times ("same"). Percentages show the observations divided by the sum of unique and redundant CSMs (rounded). The theoretical accuracy limit for hSAX and SCX was derived by summing the percentages of unique and 'red/same' CSMs (hSAX: 73%, SCX: 65%).

Table S5: Rescoring gains with different number of chromatographic dimensions.

| | level | reference | RP | SCX-RP | hSAX-RP | SCX-hSAX-RP |
|---|---|---|---|---|---|---|
| heteromeric | CSM | 724 | 902 (+1.25x) | 977 (+1.35x) | 1092 (+1.51x) | **1199 (+1.66x)** |
| heteromeric | Peptide | 507 | 619 (+1.22x) | 664 (+1.31x) | 737 (+1.45x) | **801 (+1.58x)** |
| heteromeric | Residues | 414 | 508 (+1.23x) | 546 (+1.32x) | 603 (+1.46x) | **654 (+1.58x)** |
| heteromeric | PPI | 109 | 135 (+1.24x) | 131 (+1.2x) | **157 (+1.44x)** | 152 (+1.39x) |
| self | CSM | 10357 | 10404 (+1.0x) | 10428 (+1.01x) | 10439 (+1.01x) | **10443 (+1.01x)** |
| self | Peptide | 6521 | 6565 (+1.01x) | 6586 (+1.01x) | 6598 (+1.01x) | **6601 (+1.01x)** |
| self | Residues | 4810 | 4853 (+1.01x) | 4873 (+1.01x) | 4886 (+1.02x) | **4888 (+1.02x)** |
| self | PPI | 478 | 514 (+1.08x) | 531 (+1.11x) | 540 (+1.13x) | **543 (+1.14x)** |

*Note*: The data corresponds to all *E. coli* target-target identifications at 5% CSM-, Peptide-, Residue-level FDR and 1% PPI-FDR. Rescoring was performed using a linear SVM. Highest values are marked in bold. The hyper-parameters for the rescoring were chosen dynamically via cross-validation for each run ('class_weight': 'None', all conditions; 'C': 100 (RP, SCX-RP, SCX-hSAX-RP); 'C': 10 (hSAX-RP)), according to the sklearn API. Values are rounded to two digits.

### S4 pLink2 Processing

The recalibrated MGF files were searched with pLink 2 (2.3.9) with the following search parameters: Flow Type, HCD; Cross-Linker, DSS with AlphaSites = BetaSites = [KSTY; Enzyme, Trypsin; Missed Cleavages, 2; Peptide Mass [600, 6000]; Precursor Tolerance, 5ppm; Fragment Tolerance, 3ppm; Fixed Modifications, Carbamidomethylation[C]; Variable Modifications, Oxidation[M]. The filter parameters were as follows: Filter Tolerance, 10 ppm; FDR, separate FDR 1% at CSM level; Compute E-value, False.

We further processed the unfiltered results table from pLink2 in order to get all CSMs (including decoys) and their associated error estimates for usage in xiRT. In short, we added the information about peptide origin, target-decoy origin, species, peptide positions and RT in SCX/hSAX/RP. These steps were only performed with the peptides that pass the 0.5 Q-value threshold (similar to the xiSEARCH processing, as only 50% CSM-FDR data was used). These additional annotation steps were necessary since the filtered pLink2 results (*.filtered_cross-linked-spectra) do not provide the necessary information (e.g. decoy hits and error estimates are not provided).

The generated file was then used as input for xiRT. For xiRT, the same settings as for xiSEARCH were used. In total 35822 peptides were used as input data. During the CV 3866 peptides were used for training, 430 for validation and 2147 for prediction (prediction-fold is visualized in Fig. S15, together with a 2D-feature space representation using UMAP[8]. Using crosslinks identified from pLink2 lead to a comparable prediction performance as using xiSEARCH (3871 training peptides, 431 validation peptides, 2151 prediction peptides).
